# Quantification of the mobility potential of antibiotic resistance genes through multiplexed ddPCR linkage analysis

**DOI:** 10.1101/2022.09.26.509453

**Authors:** Magali de la Cruz Barron, David Kneis, Alan Xavier Elena, Kenyum Bagra, Thomas U. Berendonk, Uli Klümper

## Abstract

Antibiotic resistance genes (ARGs) are widely disseminated within microbiomes of the human, animal, and environmental spheres. There is a clear need for global monitoring and risk assessment initiatives to evaluate the risks of ARGs towards human health. Therefore, not only ARG abundances within a given environment, but also their mobility, hence their ability to spread to human pathogenic bacteria needs to be quantified. Consequently, methods to accurately quantify the linkage of ARGs with mobile genetic elements are urgently needed.

We developed a novel, sequencing-independent method for assessing ARG mobility by combining multiplexed droplet digital PCR (ddPCR) on DNA sheared into short fragments with statistical analysis. This allows quantifying the physical linkage between ARGs and mobile genetic elements, here demonstrated for the sulfonamide ARG *sul1* and the Class1 integron integrase gene *intI1*. The method’s efficiency is demonstrated using mixtures of model DNA fragments with either linked and unlinked target genes: Linkage of the two target genes can be accurately quantified based on high correlation coefficients between observed and expected values (R^2^) as well as low mean absolute errors (MAE) for both target genes, *sul1* (R^2^=0.9997, MAE=0.71%, n=24) and *intI1* (R^2^=0.9991, MAE=1.14%, n=24). Furthermore, we demonstrate that the chosen fragmentation length of DNA during shearing allows adjusting the rate of false positives and false negative detection of linkage. The applicability of the developed method for environmental samples is further demonstrated by assessing the mobility of *sul1* across a wastewater treatment plant.

The presented method allows rapidly obtaining reliable results within hours. It is labor- and cost-efficient and does not rely on sequencing technologies. Furthermore, it has a high potential to be scaled up to multiple targets. Consequently, it merits consideration to be included within global AMR surveillance initiatives for assessing ARG mobility.

**Author Abstract:** Antibiotic resistance represents a major problem in treating bacterial infections and endangers public health. Antibiotic resistant genes (ARGs) can spread among microbes in humans, animals, and the environment, thanks to mobile genetic elements. These are genetic structures involved in the mobility of genetic information and hence microbial traits. Methods that allow simultaneously quantifying the abundance of ARGs together with their association/linkage with mobile genetic elements are fundamental for assessing the risk they pose to human health, as mobility increases their likelihood to spread to human pathogens. Here we developed a novel method that allows to quantify the abundance and the linkage between ARGs and mobile genetic elements. The method relies on droplet digital PCR (ddPCR) technology performed on fragmented environmental DNA (of a chosen size), combined with statistical analysis. We found that the method accurately quantifies the linkage between the two targets in model and environmental DNA. The method is delivering rapid, labor- and cost-efficient results as it does not rely on technology that require prior bioinformatics knowledge and can be included in future monitoring frameworks.

## Introduction

The rise in antibiotic resistance in bacterial pathogens over time represents one of the biggest challenges in treating bacterial infections [1]. Horizontal gene transfer, facilitated by mobile genetic elements (MGEs), has contributed significantly to the spread of antibiotic resistance genes (ARGs) among different hosts, accelerating this global health problem [2,3]. Addressing the problems associated with antibiotic resistance, requires action across the human, animal and environmental spheres within a “One Health” concept [4]. Within this context, there is a clear need for global monitoring and risk assessment initiatives to assess the large-scale dissemination of ARGs across human, animal, but also the environmental spheres [5]. In the latter, this proves especially difficult due to the high complexity of gene transfer and the diversity of microbial communities [6,7].

The occurrence and distribution of antimicrobial resistant bacteria (ARB), ARGs and MGEs in different environments has been extensively documented using diverse approaches ranging from selective cultivation, qPCR of ARGs and mobility related genes and metagenomic sequencing [8–11]. However, the usual report of absolute and relative abundance of ARGs is not enough to correctly assess the risks of these markers on human health, as ARGs can be found in a variety of different hosts and on diverse mobile genetic elements. The genetic and host context in which these genes appear can result in varying risk levels.

Therefore, the urgent need for a conclusive framework addressing this issue in environmental samples is beyond dispute [2,6,7]. Recently, Zhang et al. [12] proposed four key indicators to evaluate the health risk of environmental ARGs: a) Human accessibility: Is the ARG shared between human and environmental microbiomes?; b) Human pathogenicity: Has the ARG been found in human pathogens?; c) Clinical relevance: Has the presence of the ARG been associated to clinical complications/worsened clinical outcome?; d) Mobility: Is the ARG encoded on a mobile genetic element that allows transferability among bacteria, especially from non-pathogenic to pathogenic ones? The first three criteria are reasonably easy to assess with the currently available data and methods. However, evaluating the linkage of these ARGs to bacterial hosts and especially MGEs, hence their mobility in environmental communities, is crucial but remains a challenge.

Different approaches or combinations of methods have been used to link ARGs to MGEs in non-clinical environments [13]. One of the most commonly applied methods is metagenomic assembly of genomes or contigs [14–16]. However, for this approach assembled contigs have to be long enough to contain ARGs as well as genetic indicators of the mobile genetic elements in question. Further, the relative abundance of ARGs in most natural environments is low, which increases the requirements regarding sequencing depth and hence the costs [17]. Other approaches use cross-linking of chromosomal and plasmid DNA previous to long-read sequencing (Hi-C) to determine the linkage of MGEs, e.g. plasmid with associated ARGs, with their hosts [18]. However, this method is prone to wrong associations of ARGs due to methodological artifacts when cells cluster together [18] and equally suffers from the low relative abundance of mobile ARGs in most environments. Emulsion paired isolation and concatenation PCR (epicPCR) which involves the encapsulation of individual cells and fusion of ARGs to phylogenetic markers during the PCR ahead of subsequent sequencing is another method for linking target ARGs with their hosts and MGEs [19,20], but chromosomal ARGs could wrongfully be associated with MGEs co-occurring within the same bacterial cell.

The most promising approach currently available to assess ARG mobility is Inverse PCR of the ARG in question combined with long read sequencing of the amplified surrounding regions [21] to identify clusters in which ARG are co-occurring with genes associated with MGEs. This provides a comprehensive overview of the genetic environments an ARG occurs in in a given sample or environment, but it does not provide an accurate quantitative assessment of ARG mobility relative to their abundance.

Despite their individual weaknesses, these methods can provide promising insights into ARG mobility potential in the environment in the future. Still, the above mentioned approaches are technologically complicated and rely on sequencing analysis which is expensive, time consuming, and requires specialized bioinformatic knowledge for data analysis. Further, they are mainly qualitative or semi-quantitative at best, which makes risk assessments based on ARG mobility difficult. Consequently, to be included in a future global, environmental AMR monitoring framework, methods for quantitatively assessing ARG mobility that are easier applicable and of lower costs are urgently needed. Here, such a quantitative method has been developed by taking advantage of multiplexed droplet digital PCR (ddPCR) technology.

The ddPCR technology is a binary endpoint measurement based on the random distribution of nucleic acid molecules extracted from a sample partitioned into thousands of volumetrically defined droplets within a water–oil emulsion with PCR reactions taking place in each individual droplet [22]. Using a fluorescent probe based approach with different fluorochromes for different target genes allows multiplexing their detection within the entire sample, but also within individual droplets [23]. Since individual DNA molecules are randomly encapsulated within the droplets based on the Poisson distribution [24], the physical linkage between a target ARG and a target MGE on this DNA molecule can be detected and estimated by ddPCR linkage analysis using multiplexed assays in combination with statistical analysis.

Here we developed a new duplex ddPCR approach to quantify an ARGs mobility, thus the percentage of this ARGs detected copies in direct physical linkage with a marker gene associated with mobile genetic elements. This is here presented for the sulfonamide resistance gene *sul1* and its linkage to class 1 integron mobile genetic element through its integrase gene *intI1*. Although these two genes can occur individually in different environments and bacteria, they have been regularly described to co-occur in mobile integron cassettes in a wide variety of bacteria and environments [25], and hence provide a perfect study system to demonstrate the validity of the proposed method.

To prove that multiplexed ddPCR can be used as a suitable, rapid, and less expensive approach to determine the linkage between ARGs and MGEs, model DNA mixtures with linkages ranging from 100% to 0% were analyzed. The validity of the linkage detection for each of the two genes was then assessed by comparing the linkage determined for each of the mixtures using the statistical evaluation of the duplex ddPCR assay with the theoretical, expected linkage percentage. To avoid a biased linkage detection when the two target genes occur individually (outside of an individual integron cassette) on the same chromosome, extracted DNA needs, prior to analysis, to be sheared into short fragments. Based on the statistical probability evaluation of false positive detections, a shearing size of around 20,000 base pairs (20 kbp), approximately five to ten times the size of an average class 1 integron cassette [26], is suggested for optimal detection of the association of the ARG with class 1 integrons. Using the presented multiplexed ddPCR approach we demonstrate its suitability by simultaneously assessing the abundance of the individual targets and their linkages in environmental samples from the influent and effluent of a wastewater treatment plant.

## Material & Methods

### Linked and unlinked target DNA fragments

To prove the concept and optimize the methodology for determining the physical linkage between an antibiotic resistance gene and a mobility marker using a ddPCR multiplexed protocol the sulfonamide resistance genes *sul1* and the class 1 integron integrase gene *intI1* were selected. These two genes have been regularly described to co-occur in mobile integron cassettes in a wide variety of bacteria and environments [25], but can also occur independently in the same bacterial host on chromosomes or plasmids. To analyze, if the linkage of these two genes can be correctly predicted using the ddPCR protocol, we used model DNA molecules on which either one of the two targets, or both targets in linked form on the same molecule, are present.

The pNORM plasmid (http://www.norman-network.net/) served as the linked control, as it contains a single copy of both target genes in close proximity of 250 bp [9]. Plasmid DNA was extracted from the *E. coli* host strain using the Monarch® Plasmid Miniprep Kit (NEB, Ipswich, MA, USA). The extracted plasmid was linearized with restriction enzyme *BamHI* (Promega, Madison, WI, USA), and thereafter purified with QIAquick PCR Purification Kit (Qiagen, Hilden, Germany). All steps were carried out according to the protocols supplied by the manufacturers.

Purified PCR products of the *sul1* and *intI1* gene were used as the no linkage controls. PCR was carried out individually for each gene using GoTaq® Green Master Mix (Promega), with primers listed in Table 1, and linearized pNORM DNA as the template. The PCR program was set as follows: 95 °C for 10 min followed by 35 cycles at 94 °C for 30 s, 60 °C for 30 s, 72 °C for 30 s, and a final elongation cycle at 72 °C for 10 min. Successful amplification and correct size of the *sul1* and *intI1* PCR produces were finally confirmed by 1.5% agarose gel electrophoresis (40 min, 110V) and subsequently purified with the QIAquick PCR Purification Kit (Qiagen) according to the manufacturer’s instructions.

**Table 1.**
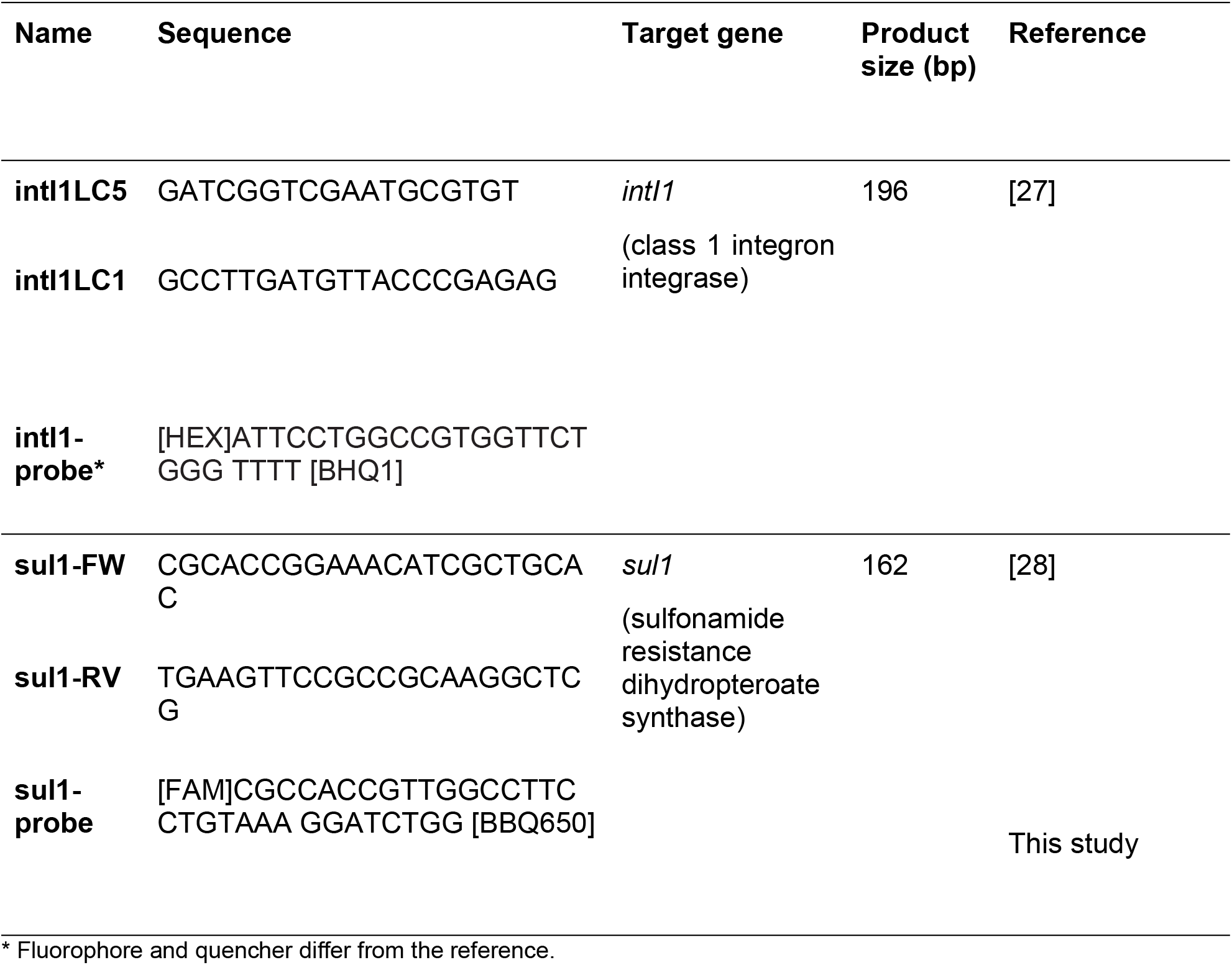
Primers and probes used in this study.

Linearized pNORM with both targets linked as well as a 1:1 mixture of the individual, unlinked PCR products were prepared in DNAse free H_2_O (Qiagen) and adjusted to 1 ng μL^-1^. These two solutions were then used in mixtures of varying ratios to assess the detection of different linkage percentages (see below) using the ddPCR duplex detection protocol.

### ddPCR duplex detection protocol

To allow the simultaneous detection of two genes within one ddPCR assay, individual probe based PCR assays for each of the genes are necessary. Each assay consists of forward and reverse primer and a probe labeled with non-interfering fluorochromes. While for the *intI1* gene such a complete assay already existed [27], for *sul1* an existing and well-validated qPCR primer set was used [28]. The matching fluorescent probe was designed in this study and validated for ddPCR detection of the *sul1* gene (Table 1). No cross-reactivity between the assays was detected when multiplexing the assays with the *intI1* probe labeled with HEX and the *sul1* probe labeled with FAM reporter fluorophores.

To quantify the two target genes *sul1* and *intI1*, we performed a duplexed ddPCR assays using the primers and probes listed on Table SI1, and the QX200 Droplet Digital PCR System (Bio-Rad, Hercules, CA, USA). Reactions were prepared in 20 μL volume, containing: ddPCR Supermix for Probes (No dUTP, Bio-Rad), each of the four primers, and two probes (Table 1) at final concentrations of 900 nm for primers and 250 nM for probes, and 1 ng of DNA. This DNA could either be a mixture of the linked and non-linked targets created previously, or genomic DNA from sheared or non-sheared samples (see below).

Droplets were generated on a QX200 droplet generator (Bio-Rad), and transferred into a 96-well PCR plate (heat-sealed with a foil plate seal, Bio-Rad). PCR was carried out in a C1000 thermal cycler (Bio-Rad) using the following cycling conditions: 95 °C for 10 min followed by 45 cycles at 94 °C for 30 s, and 60 °C for 90 s, and a final cycle at 98 °C for 10 min. A ramp rate was set to 2.0°C/s. The droplets were read using a QX200 droplet reader and data was analyzed using QuantaSoft Software v1.4.0.99 (Bio-Rad).

#### Calculation of the linkage of genes

During the ddPCR partitioning process, DNA fragments are randomly Poisson distributed into thousands of droplets (∼20,000 per ddPCR assay) [22]. This means that individual droplets could either be empty or contain one or more DNA fragments with or without target(s) of interest (Fig. 1 A). In each of these droplets the PCR reaction is performed and the fluorescence of each droplet detected. After detection and data analysis, in multiplex ddPCR experiments like performed here, four clusters should be observed, which are orthogonal to each other (Fig. 1 B). Each droplet could either contain no target, only target A (*sul1*), only target B (*intI1*), or both targets either by chance (co-localization) or due to physical linkage on the same DNA fragment (Fig 1 B).

**Fig. 1.**
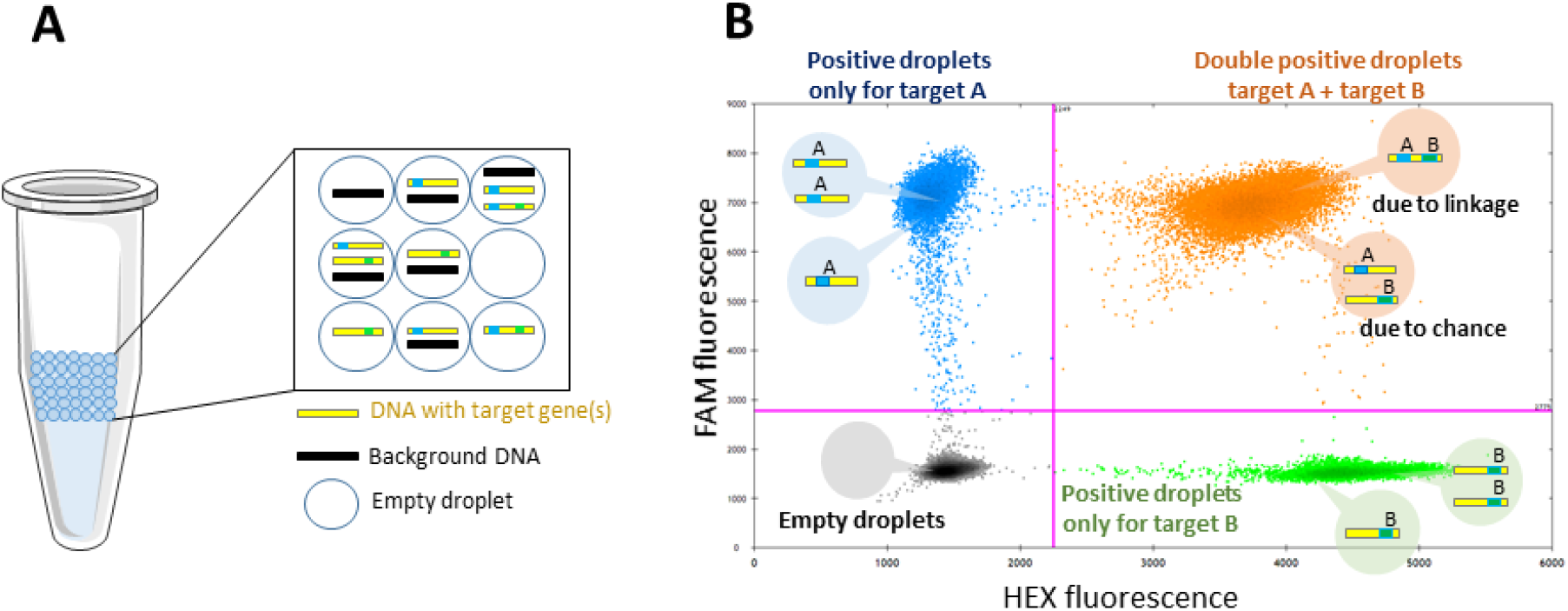
(A) Partitioning of PCR reaction into ca. 20,000 droplets of uniform size and volume, containing target and background DNA. (B) Data output from multiplex ddPCR experiments. Droplets form 4 clusters arranged orthogonally to each other. In grey: Empty droplets, negative for both targets (- -); Blue: positive droplets only for target A (+ -); Green: positive droplets only for target B (- +) and Orange: double-positive droplets (+ +) due to chance (no linkage) or due to physical linkage.

Total concentrations of either target A or target B can simply be calculated based on the Poisson distribution and the amount of droplets positive for the specific target. The total number of physically linked molecules can then be calculated by determining the excess of double-positive droplets over that expected due to chance. To determine this linkage between the two target genes mathematically a modified version of the equations described by Regan et al. [24] for eukaryotic chromosomal linkage analysis needs to be applied:

If the two markers A and B are unlinked, their presence in individual droplets follows the Poisson distribution. This includes the possibility of multiple DNA fragments each containing copies of the same target or each containing individual copies of both targets appearing in a single droplet by chance. Consequently, they will be partitioned according to the following equation, with N as the number of droplets analyzed from a ddPCR assay, N_A_ and N_B_ the number of droplets for the single target, N_E_ the number of empty, double-negative droplets and N_AB_ the number of double positive droplets:

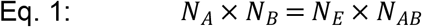

Rearranging this equation allows calculating the amount of double-positive droplets that appear purely by chance N_ch_ of both unlinked targets appearing in the same droplet:

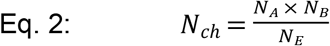

When target A and B are physically linked (<u>AB</u>) additional double-positive droplets that are not based on chance appear. The amount of droplets that do not contain these linked targets N_no<u>tAB</u>_ can now be calculated as the sum of empty, single-positive and chance-based double-positive droplets.

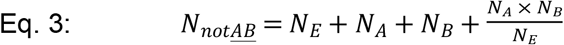

By resolving the Poisson distribution through the natural logarithm and subtracting N_no<u>tAB</u>_ from the total number of measured droplets N_tot_ we calculate the concentration (average copies / droplet) of molecules with linked targets λ<u>_AB_</u> using:

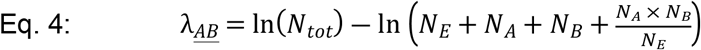

Similarly, the concentration of each individual target (λ_A_, λ_B_), irrespective of linkage, can be calculated as:

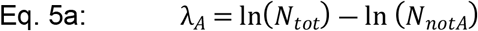

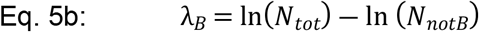

For each individual target the percentage of linked targets among the total concentration can now be calculated as:

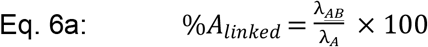

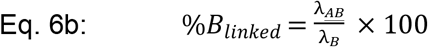

#### Proof of concept

To prove that the percentage of linked targets can be correctly estimated using the methodology described above, different voluminal ratios of the linked and unlinked target solutions were mixed at volumes (V) of linked targets at 100%, 90%, 75%, 50%, 25%, 10%, 5% and 0%. From the 100% and 0% mixtures the absolute concentration (c) of each gene in the linked and the unlinked samples were determined by ddPCR according to the protocol above. These were then used to calculate the theoretical ratio of linked to unlinked target genes in the mixtures:

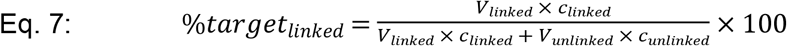

The validity of the linkage detection for each of the two genes was then assessed by comparing this theoretical linkage percentage with the value determined for each of the mixtures using the statistical evaluation of the duplex ddPCR assay.

### Environmental samples

Environmental wastewater samples of the influent and effluent of the wastewater treatment plant in Dresden-Kaditz Germany (51.072N, 13.678E) were collected in sterile flasks, and stored at 4 ºC until DNA extraction for a maximum of 12 h. Three replicate grab samples of 1L each were taken for the influent and effluent on 24th of March 2022. From each of the samples biomass was collected by filtration from a total volume of 130 mL for influent and 800 mL for effluent samples. Total environmental DNA was extracted from the filters using the DNeasy PowerWater kit (Qiagen) following the manufacturer’s instructions. DNA concentration and purity were estimated spectrophotometrically. All DNA was stored at -20°C until use for ddPCR analysis.

### DNA shearing

The two target genes can appear on one and the same DNA fragment, such as a bacterial chromosome, either linked in an individual integron cassette or independent of one another at random locations. To allow reliable analysis of their linkage within the same integron cassette, the DNA needs to be sheared into fragments of a specific size ahead of the analysis (Fig. 2).

**Fig. 2.**
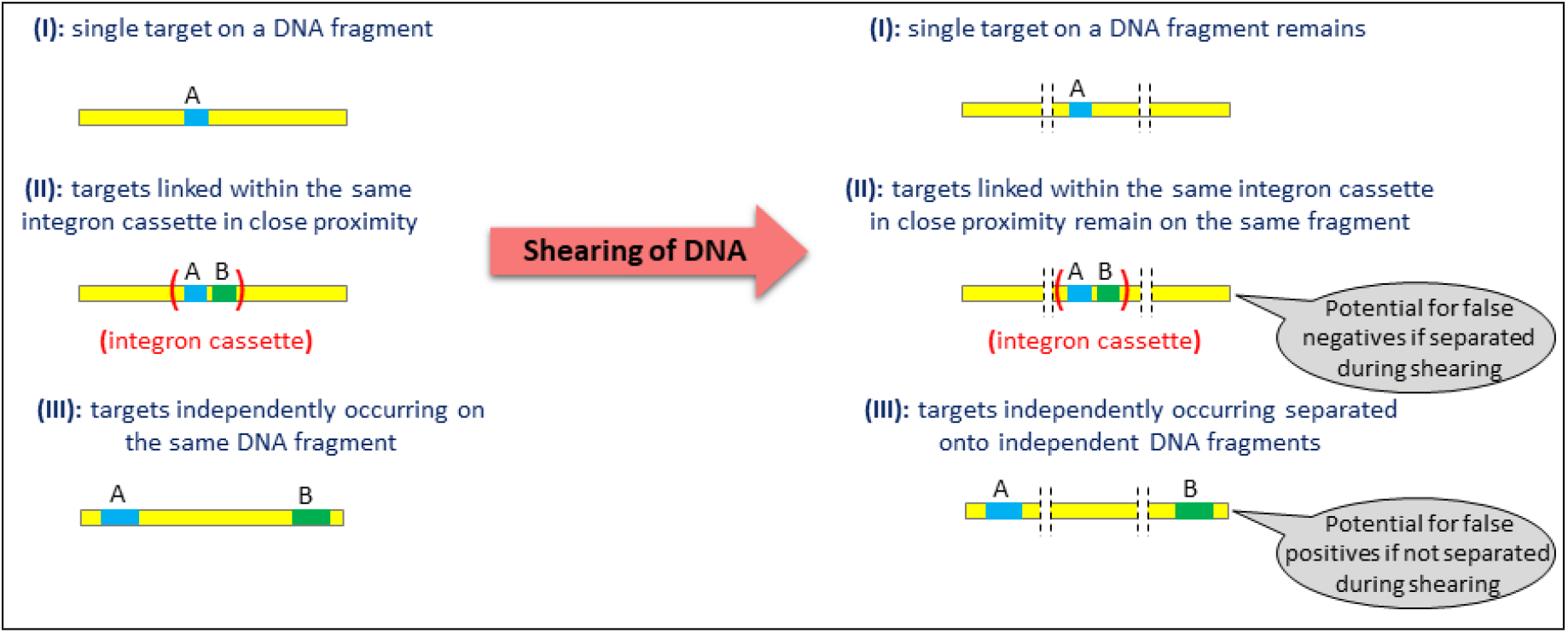
Shearing of DNA into fragments of a specific sizes allows distinguishing between both targets (A & B) occurring on one and the same DNA fragment due to either linkage within an integron cassette or both genes appearing independently. Fragmentation size allows adjusting the rate of false positive and false negative detection of linkage.

The size of the fragments generated during the process of DNA shearing crucially affects the probabilities of false positive (*P*_FP_) and false negative outcomes (*P*_FN_). As a false positive outcome we regard cases where the two different target genes are detected on one and the same DNA fragment even though the targets are not linked (e.g. they are not part of a common mobile gene cassette). Conversely, false negative refers to the failure of detecting actual linkage between the targets (e.g. by being embedded on a common gene cassette) in response to the DNA fragmentation during the shearing process.

As long as the target sequence is short in relation to the fragment length (L_F_), *P*_FP_ can be approximated as the probability of jointly detecting the two unlinked targets in any DNA fragment obtained upon shearing of a genome. Thus, if the total length of the genome is denoted L_G_, *P*_FP_ can be approximated by Eq. 8.

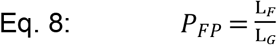

The probability of false negative results is not only a function of L_F_, but it also depends on the length of the hypothetical gene cassette (LC) harboring the two targets and the actual position of the targets within the cassette. If we focus on cases where L_C_ < L_F_, i.e. a cassette can be sheared into no more than two fragments and the two targets are positioned in maximum distance within the cassette (representing the worst case scenario), *P*_FN_ can be approximated by Eq. 9.

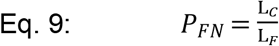

Estimates of *P*_FP_ and worst-case estimates of *P*_FN_ are reported for a range of assumed fragment lengths (Table 2). For these simulations, based on the literature, we assumed an average bacterial chromosome size L_G_ of 5 Mbp [29] and the maximal length of clinically relevant Class1 integron cassettes for L_C_ of 5 kbp [26]. This provides insights into the worst case scenario of the genes being linked but occurring in the same cassette at a maximal distance of 5 kbp. Since the two types of error show opposing dependencies on the fragment length, DNA shearing needs to target at intermediate values of L_F_ such that *P*_FP_ and *P*_FN_ are in balance. Under the given scenario, reasonable target values of L_F_ are in the range of 20 to 50 kbp.

**Table 2.** Probabilities of false positive (*P*_FP_, Eq. 8) and false negative outcomes (*P*_FN_, Eq. 9) in relation to assumed fragment lengths (L_F_) resulting from DNA shearing. Reported numbers refer to a scenario where L_G_=5 Mbp (total length of genome) and L_C_=5 kbp (length of gene cassette). Note that the values reported for P_FN_ represent worst-case estimates.

| $L_F$ (kbp) | $P_{FP}$ | $P_{FN}$ |
| --- | --- | --- |
| 10 | 0.002 | 0.5 |
| 20 | 0.004 | 0.25 |
| 50 | 0.01 | 0.1 |
| 100 | 0.02 | 0.05 |

Accordingly, a fragment length of 20 kbp was chosen for this proof of concept study to achieve a rate of false positive detection of less than 0.4% of the cases. The Covaris® g-TUBE (Covaris, Woburn, MA, USA) was used to shear the genomic DNA into the selected fragment sizes of around 20 kb, according to the protocol provided by the manufacturer. Success of the shearing process into fragments of the desired length was confirmed using gel electrophoresis.

## Results

### Reliable detection of different levels of linkage between the two target genes *sul1* and *intI1* from model samples

To determine if linkage between *sul1* and *intI1* is detectable using the statistical evaluation of the proposed multiplexed ddPCR method, initially the plasmid on which the two targets are linked and the unlinked PCR products of the two targets were used as templates. In the linked control sample, the plasmid DNA molecule hosting the linked copies of the two target genes was added at approximately 1,500 copies μL^-1^. The concentrations of the two target genes measured using the multiplexed ddPCR protocol was 1,510.0 ± 26.9 (mean ± SD) copies μL^-1^ for *sul1* and 1,448.0 ± 20.0 copies μL^-1^ for *intI1*. This accounted for an acceptable average detection efficiency of 100.7 % and 96.5 % and further confirmed that both genes indeed appeared in a single copy on the used model plasmid.

Obtained linkage percentages were calculated from the distribution of droplets (Fig. 3 A) after correcting for stochastic effects (see M&M for calculation) as 92.42 ± 0.62 % (*sul1*) and 96.34 ± 0.19 % (*intI1*). While significantly different from the expected linkage percentage of 100% (both *p*<0.05, two-tailed *t*-test, dF=5), the rate of false negatives, and hence the average underestimation of gene linkage of 7.58% and 3.66% for the two genes (rate of false negatives) falls still far below the worst case scenario calculated for the optimization of the shearing fragment length of a maximal 25 % (Table 2).

**Fig. 3.**
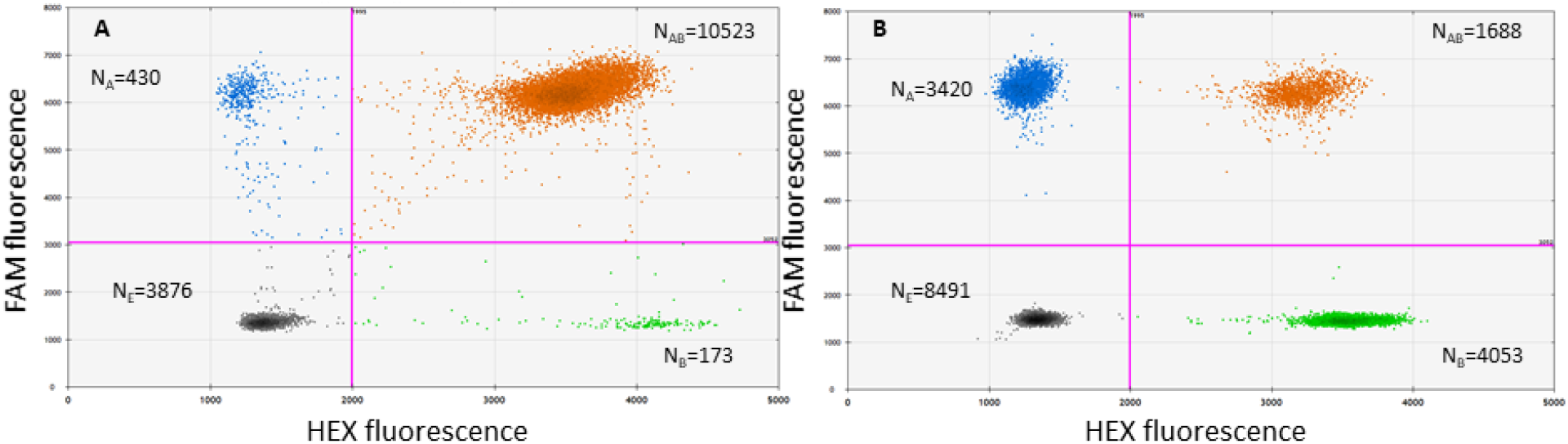
Data output from ddPCR experiments using the 100% (A) and 0% (B) linkage controls. N is the number of accepted droplets analyzed. Grey: Empty droplets (N_E_), negative for both targets; Blue: droplets positive only for target A, here *sul1* (N_A_); Green: droplets positive only for target B, here *intI1* (N_B_); Orange: droplets double-positive for both targets including those due to physical linkage and due to chance (N_AB_).

In the unlinked control still around 5% of droplets that contained both targets were obtained (Fig. 3 B). Statistical evaluation of the results revealed, however, that these occurred exclusively by chance with the obtained linkage percentages for the two target genes calculated as 0.31 ± 1.05 % (*sul1*) and 0.26 ± 0.92 % (*intI1*). These results were not significantly different from the expected linkage of 0% for either of the genes (*p*_*sul1*_=0.658, *t*_*sul1*_=0.515; *p*_*intI1*_=0.663, *t*_*intI1*_=0.506; two-tailed *t*-test, dF=5), hence demonstrating that if targets are not linked, no false positives for their potential association on the same DNA molecule can be detected.

When validating the method by using mixtures of linked and unlinked targets at defined linkage percentages the experimentally obtained values were very well correlated with the theoretically expected ones based on the correlation coefficient and the mean absolute error for the linkage of both target genes, *sul1* (R^2^=0.9997, MAE=0.71%, n=24) and *intI1* (R^2^=0.9991, MAE=1.14%, n=24) (Fig. 4). Further the method proved robust as the standard deviation of the technical replicate measurements at different linkage ratios was consistently low (0.18-1.08%).

**Fig. 4.**
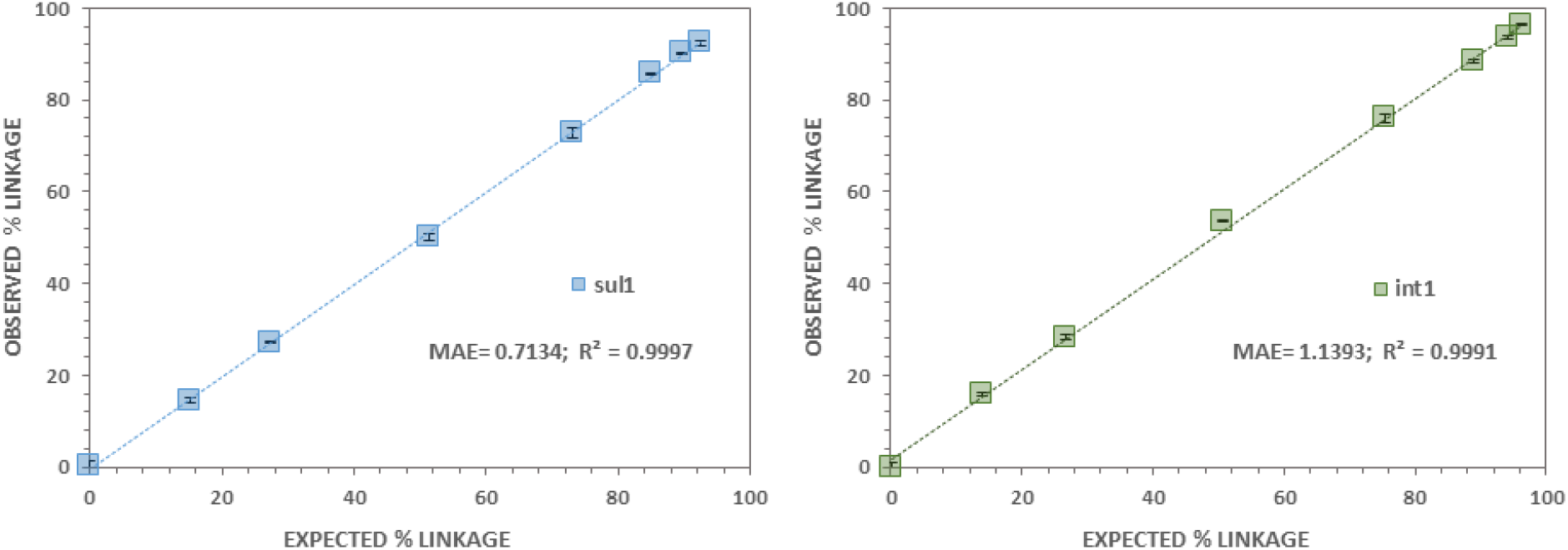
Linear correlation between theoretical and experimentally calculated linkage percentage for the two target genes *sul1* (A) and *intI1* (B). Tested samples were prepared at eight defined volumetric mixtures of linked and unliked targets and measured with 3 technical replicates by multiplexed ddPCR (n=24). MAE, mean absolute error in percentage (%).

Consequently, the proposed method provides reliable values for the linkage between the target genes, with the level of false positive detection of linkage being negligible and the level of false negative detection of linkage at high linkage percentages being within the level expected due to methodological constraints based on the optimization of the shearing procedure. This allowed applying the method to more complex environmental samples in order to detect the mobility and transferability potential of the *sul1* gene through linkage with class1 integrons.

### Abundance of target genes in wastewater influent and effluent samples

The optimized protocol was applied to environmental water samples obtained from the influent and effluent of a wastewater treatment plant to first determine the abundance of the target genes. The abundance of both target genes *sul1* and *intI1* was significantly higher in the influent compared to the effluent of the wastewater treatment plant (*p*_*sul1*_<0.0001, *t*_*sul1*_=36.72; *p*_*intI1*_<0.0001, t_*intI1*_=41.18; dF=10, two-tailed *t*-test). In the influent the abundance of *sul1* was determined as 9.15 ± 0.60 × 10^5^ copies mL^-1^ and the abundance of *intI1* was 1.58 ± 0.09 × 10^6^ copies mL^-1^, while in the effluent these numbers were reduced by 58-fold to 1.56 ± 0.22 × 10^4^ copies mL^-1^ (*sul1*) and 64-fold to 2.49 ± 0.34 × 10^4^ copies mL^-1^ (*intI1*) (Fig. 5 A).

**Fig. 5.**
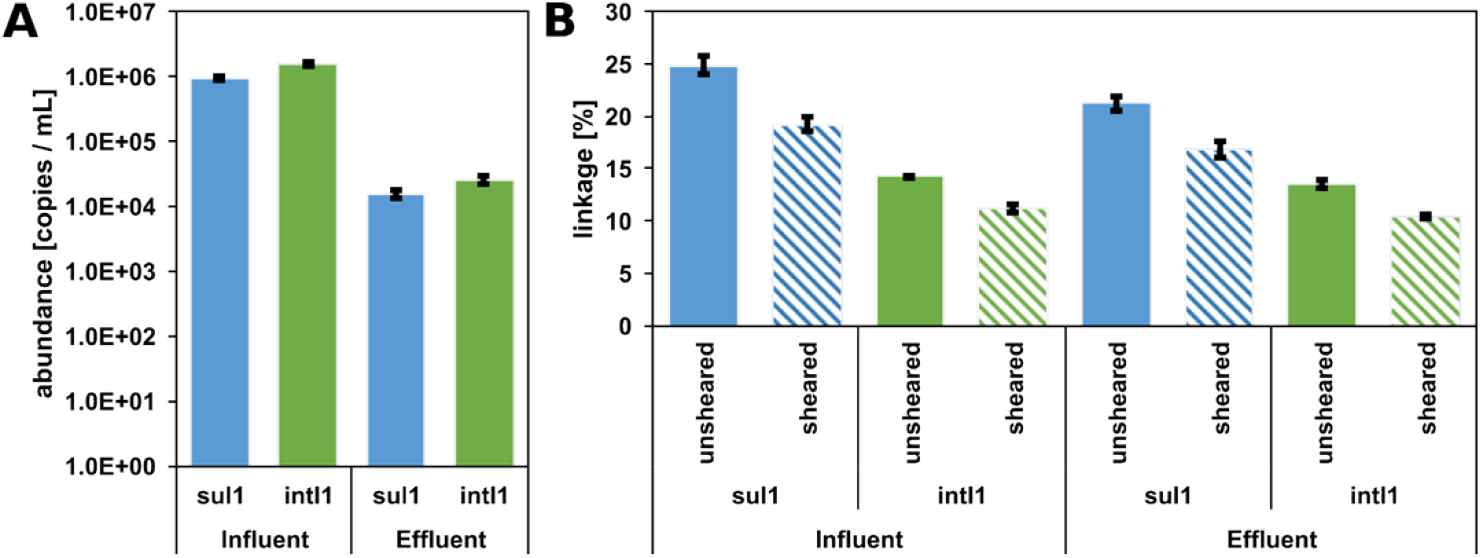
Abundance (A) and linkage (B) of the two target genes *sul1* and *intI1* in wastewater influent and effluent samples. Linkage was determined for unsheared as well as sheared DNA (average fragment size = 20 kbp).

### Linkage between the *sul1* and *intI1* target genes in unsheared DNA of the environmental samples

Based on the statistical evaluation of the multiplexed ddPCR assay, the detected initial linkage when applying the unsheared DNA, extracted using the commercial kit, was also reduced when comparing the influent with the effluent. For *sul1* the percentage of detected gene copies that were linked to *intI1* was 24.83 ± 0.88 % in the influent and significantly reduced to 21.25 ± 0.71 % in the effluent (*p*=0.0054, *t*=5.47, dF=4, *t-*test, Fig. 5 B). Equally for *intI1* the percentage of gene copies linked to *sul1* was significantly lowered during passage through the wastewater treatment plant from 14.21 ± 0.17 to 13.53 ± 0.30 % (*p*=0.0278, *t*=3.38, dF=4, *t*-test, Fig. 5 B). The significantly lower linkage of *intI1* compared to *sul1* in all samples is here due to the significantly higher abundance of *intI1* in both sample types. However, the unsheared DNA does not account for the co-abundance of the two genes on longer DNA fragments irrespective of co-occurrence within the context of a class 1 integron cassette.

### Shearing of DNA reveals the true linkage between the *sul1* and *intI1* target genes in environmental samples

Based on the calculations laid out in the Materials and Methods section, DNA was subsequently sheared using the Covaris® g-TUBE to the optimal fragment size of ∼20 kbp. Shearing did not have an effect on the detection of either of the two target genes as the ratio of *sul1* to *intI1* between the unsheared (Influent: 0.57 ± 0.02; Effluent: 0.64 ± 0.01) and sheared assays (Influent: 0.58 ± 0.00; Effluent: 0.61 ± 0.02) remained insignificantly affected (*p*_Inf_=0.4588, *t*_Inf_=0.819; *p*_Eff_=0.1516, *t*_Eff_=1.769; dF=4, two-tailed *t*-test). However, shearing had a significant effect in reducing the detected linkages of each of the genes for each sample type (Fig. 5 B; all *p*=0.0001-0.0017, dF=4, two-tailed *t*-test). For *sul1* the linkage to *intI1* after shearing was reduced to 19.25 ± 0.62 % in the influent and 16.87 ± 0.73 % in the effluent. For *intI1* the linkage after shearing was determined as 11.24 ± 0.43 % in the influent and 10.36 ± 0.18 % in the effluent.

Taking these new linkage values into account, the wastewater treatment process did not only reduce the gene abundance of the two target genes, but also significantly reduced their linkage and hence the mobility of the *sul1* ARG (*p*=0.0125, *t*=4.31, dF=4, two-tailed *t*-test) (Fig. 5 B). However, this reduction was only very slight compared to the 10-to 100-fold reduction in gene absolute abundance per mL.

Overall, this shows that the method can be easily applied for mobility analysis of ARGs within diverse environmental samples. Based on the results obtained without DNA shearing, the linkage between the two genes would have been overestimated by around 20% for each of the genes in each of the samples, due to independent occurrence of both targets (outside of the identical integron casette) on the same DNA fragment. Thus, DNA fragmentation is an absolute necessity for an accurate detection of ARG mobility.

## Discussion

We here present a novel, multiplexed ddPCR based approach to simultaneously assess the abundance of individual target genes and their linkage from complex environmental samples. This allows estimating the linkage of ARGs to mobile genetic elements, hence their mobility within a given sample. This is an important contributor to the risk an environmental ARG is posing towards human health as it describes the likelihood of an ARG being able to spread to a human pathogen [12]. This is exemplarily shown for the sulfonamide resistance gene *sul1* and its association with the class 1 integron cassettes based on the integron integrase gene *intI1*. Using model DNA target molecules with either each gene in isolation or both genes linked, we demonstrate that the theoretical linkage in mixtures of these targets can indeed be accurately quantified. Further, the applicability to assess such linkages in environmental samples is demonstrated based on wastewater influent and effluent samples.

For integron cassettes that can equally be part of chromosomes or other mobile genetic elements [30] we, through means of statistical modeling, demonstrate that the rates of false positive and false negative detections depend directly on the shearing fragment length of the sample DNA. In this proof of principle study, a conservative fragment length of 20 kbp for class 1 integron cassettes of an average size of 5 kbp [26] was chosen to keep the rates of false positives below 0.4% based on the mathematical assessment of the worst case scenario of both genes being on opposite sides of the integron cassette. This cutoff comes with a tradeoff of a relatively high 25% theoretical maximal false negative detection rate. Using our reference plasmid with linked targets in close proximity the experimentally detected rate of false negative detection of linkage was however far lower at between 3 and 8 %, hence indicating that optimizing fragment length for low rates of false positive detection might be favorable.

We further prove that DNA fragmentation is an absolute necessity for an accurate detection of ARG mobility in environmental samples as the linkage between the two genes in the wastewater samples is overestimated by around 20% in the absence of the fragmentation process. Obviously when applying this method in the future, different fragment sizes for different combinations of targets lead to different tradeoffs between false positive and false negative detection and need to be carefully decided based on the task at hand.

Currently the main methods for evaluating the linkage of ARGs with mobile genetic elements is the use of metagenomic sequencing with contig assembly and annotation [e.g. 15] or the use of Inverse PCR combined with long read sequencing [21]. In metagenomic sequencing, due to sequencing depth constraints, only the most common ARGs with a relative abundance of more than 10^−4^ copies per 16S copy can be detected [31]. PCR based approaches, including Inverse PCR and the here presented ddPCR approach, have several orders of magnitude improved detection limits [32] making them also suitable for those ARGs that are rare in a given environment. However, while in metagenomics all ARGs and MGEs within the detection limit will be sequenced and analyzed simultaneously, PCR based approaches have the disadvantage of needing specific primers for each individual specific target. Consequently, novel ARGs or those ARGs not expected in a certain sample might be missed as they might not be tested for [17]. Inverse PCR combined with long read sequencing provides a snapshot of all MGEs a specific ARG is associated with in a given sample, necessary information in assessing an ARGs mobility potential [21]. Still this approach is also relying on sequencing. Furthermore, due to the PCR amplification steps involved, it does not allow for a quantitative analysis of mobility. If such associations with mobile genetic elements are common or rare, remains however crucial information needed in risk assessment. A major advantage of the here presented ddPCR approach lies in the cheap, rapid and straightforward generation of quantitative results. While sequencing and subsequent sequence analysis take time, computing power and expert bioinformatics knowledge, the ddPCR approach allows the generation of results within hours and the evaluation can be automated using simple scripts for statistical analysis.

In addition, the presented method has a high potential to be up-scaled to multiple targets. Currently, ddPCR machines are able to detect up to two fluorochromes simultaneously, with future developments expected to increase this number. Furthermore, it can straightforwardly be extended to other ARGs or genes of interests and different groups of mobile genetic elements. The only necessary means to achieve this is the design of appropriate sets of primers and probes that allow multiplexed ddPCR analysis for the respective targets. For example, the association of a given ARG with different groups of plasmids in an environmental sample can easily be tested. In that case the shearing process could even be omitted, as plasmids already provide separate DNA entities from chromosomes. Hence the rate of false positives would be dramatically reduced, which allows for the detection of ARG-plasmid associations directly from extracted environmental DNA. Here the presented method would benefit from a combination with Inverse PCR with long read sequencing [21] which allows determining the most interesting ARG-MGE associations, which could subsequently be quantified using the multiplexed ddPCR approach with targeted primers. Different degenerate primers for different groups of plasmids even already exist from replicon- or MOB-typing of plasmids [33–35] and could be adapted for this approach. Detection of ARG-plasmid associations could hence be carried out for individual plasmid groups using different fluorophores. In addition the use of already existing degenerate primers [34] or mixtures of primers targeting multiple plasmid groups that all use the identical fluorophores could provide a general percentage of ARG-plasmid linkage in a given sample. Such approaches could be especially valuable considering the possibility of combining them with microfluidic enrichment of ddPCR droplets based on positive fluorescence signals for targeted sequencing of those plasmids that do contain environmentally relevant ARGs [36].

In summary, the presented, multiplexed ddPCR approach proves to be a suitable and rapid, and cost-efficient approach to simultaneously assess the abundance of individual target genes and their linkage. With a straightforward protocol, high adaptability towards new targets and potential for extension through additional methods, it is hence a promising method for the assessment of ARG mobility. The suitability of ddPCR applications in environmental surveillance protocols has already been demonstrated as it has recently gained traction as a cost-efficient method included in global wastewater monitoring initiatives of the COVID-19 pandemic [37–39]. With ARG mobility as one of the main determinants for ARG risks towards human health [12], this targeted approach might hence be a suitable addition to be considered for future, global environmental AMR monitoring frameworks to assess ARG mobility.

## Acknowledgements

We would like to express our gratitude to Volker Kühn and Gerold Fritsche for access to the influent and effluent wastewater samples. We thank Christiane Zschornack for laboratory support.

## Author contributions

**Magali de la Cruz Barron**: Conceptualization; Data curation; Formal analysis; Investigation; Methodology; Validation; Visualization; Writing - original draft; Writing - review & editing.

**David Kneis**: Data curation; Formal analysis; Methodology; Visualization; Writing - review & editing.

**Alan Xavier Elena**: Conceptualization; Writing - review & editing

**Kenyum Bagra**: Investigation; Writing - review & editing

**Thomas U. Berendonk**: Funding acquisition; Project administration; Resources; Supervision; Writing - review & editing.

**Uli Klümper**: Conceptualization; Data curation; Formal analysis; Investigation; Methodology; Project administration; Supervision; Validation; Visualization; Writing - original draft; Writing - review & editing.

## References

1. Laxminarayan R, Duse A, Wattal C, Zaidi AKM, Wertheim HFL, Sumpradit N, et al. Antibiotic resistance—the need for global solutions. Lancet Infect Dis. 2013;13: 1057– 1098. doi:10.1016/S1473-3099(13)70318-9

2. Berendonk TU, Manaia CM, Merlin C, Fatta-Kassinos D, Cytryn E, Walsh F, et al. Tackling antibiotic resistance: the environmental framework. Nat Rev Microbiol. 2015;13: 310–317. doi:10.1038/nrmicro3439

3. Klümper U, Riber L, Dechesne A, Sannazzarro A, Hansen LH, Sørensen SJ, et al. Broad host range plasmids can invade an unexpectedly diverse fraction of a soil bacterial community. ISME J. 2015;9: 934–945. doi:10.1038/ismej.2014.191

4. Queenan K, Häsler B, Rushton J. A One Health approach to antimicrobial resistance surveillance: is there a business case for it? Int J Antimicrob Agents. 2016;48: 422–427. doi:10.1016/j.ijantimicag.2016.06.014

5. Hernando-Amado S, Coque TM, Baquero F, Martínez JL. Defining and combating antibiotic resistance from One Health and Global Health perspectives. Nat Microbiol. 2019;4: 1432– 1442. doi:10.1038/S41564-019-0503-9

6. Smalla K, Cook K, Djordjevic SP, Klümper U, Gillings M. Environmental dimensions of antibiotic resistance: assessment of basic science gaps. FEMS Microbiol Ecol. 2018;94. doi:10.1093/femsec/fiy195

7. Huijbers PMC, Flach CF, Larsson DGJ. A conceptual framework for the environmental surveillance of antibiotics and antibiotic resistance. Environ Int. 2019;130: 104880. doi:10.1016/j.envint.2019.05.074

8. Kampouris IDID, Agrawal S, Orschler L, Cacace D, Kunze S, Berendonk Tutu, et al. Antibiotic resistance gene load and irrigation intensity determine the impact of wastewater irrigation on antimicrobial resistance in the soil microbiome. Water Res. 2021;193: 116818. doi:10.1016/j.watres.2021.116818

9. Cacace D, Fatta-Kassinos D, Manaia CM, Cytryn E, Kreuzinger N, Rizzo L, et al. Antibiotic resistance genes in treated wastewater and in the receiving water bodies: A pan-European survey of urban settings. Water Res. 2019;162: 320–330. doi:10.1016/j.watres.2019.06.039

10. Pärnänen Kmm, Narciso-Da-Rocha C, Kneis D, Berendonk TU, Cacace D, Do TT, et al. Antibiotic resistance in European wastewater treatment plants mirrors the pattern of clinical antibiotic resistance prevalence. Sci Adv. 2019;5: eaau9124. doi:10.1126/sciadv.aau9124

11. Barrón MD la C, Merlin C, Guilloteau H, Montargès-Pelletier E, Bellanger X. Suspended materials in river waters differentially enrich class 1 integron- and IncP-1 plasmid-carrying bacteria in sediments. Front Microbiol. 2018;9: 1443. doi:10.3389/FMICB.2018.01443/BIBTEX

12. Zhang AN, Gaston JM, Dai CL, Zhao S, Poyet M, Groussin M, et al. An omics-based framework for assessing the health risk of antimicrobial resistance genes. Nat Commun 2021 121. 2021;12: 1–11. doi:10.1038/s41467-021-25096-3

13. Rice EW, Wang P, Smith AL, Stadler LB. Determining Hosts of Antibiotic Resistance Genes: A Review of Methodological Advances. Environ Sci Technol Lett. 2020;7: 282–291. doi:10.1021/ACS.ESTLETT.0C00202/ASSET/IMAGES/LARGE/EZ0C00202_0003.JPEG

14. Dai D, Brown C, Bürgmann H, Larsson DGJ, Nambi I, Zhang T, et al. Long-read metagenomic sequencing reveals shifts in associations of antibiotic resistance genes with mobile genetic elements from sewage to activated sludge. Microbiome. 2022;10: 1–16. doi:10.1186/S40168-02101216-5/FIGURES/6

15. Kneis D, Berendonk TU, Forslund SK, Hess S. Antibiotic Resistance Genes in River Biofilms: A Metagenomic Approach toward the Identification of Sources and Candidate Hosts. Environ Sci Technol. 2022. doi:10.1021/ACS.EST.2C00370/ASSET/IMAGES/LARGE/ES2C00370_0006.JPEG

16. Liu Z, Klümper U, Liu Y, Yang Y, Wei Q, Lin JG, et al. Metagenomic and metatranscriptomic analyses reveal activity and hosts of antibiotic resistance genes in activated sludge. Environ Int. 2019;129: 208–220. doi:10.1016/j.envint.2019.05.036

17. Klümper U, Leonard AFC, Stanton IC, Ajun MF, Antonio M, Balkhy H, et al. Towards Developing an International Environmental AMR Surveillance Strategy. In: JPIAMR Report [Internet]. 2022 [cited 5 Sep 2022]. Available: https://www.jpiamr.eu/projects/towards-developing-an-international-environmental-amr-surveillance-strategy/#popular-summary

18. Stalder T, Press MO, Sullivan S, Liachko I, Top EM. Linking the resistome and plasmidome to the microbiome. ISME J 2019 1310. 2019;13: 2437–2446. doi:10.1038/s41396-019-0446-4

19. Hultman J, Tamminen M, Pärnänen K, Cairns J, Karkman A, Virta M. Host range of antibiotic resistance genes in wastewater treatment plant influent and effluent. FEMS Microbiol Ecol. 2018;94. doi:10.1093/femsec/fiy038

20. Wei Z, Feng K, Wang Z, Zhang Y, Yang M, Zhu YG, et al. High-Throughput Single-Cell Technology Reveals the Contribution of Horizontal Gene Transfer to Typical Antibiotic Resistance Gene Dissemination in Wastewater Treatment Plants. Environ Sci Technol. 2021;55: 11824–11834. doi:10.1021/ACS.EST.1C01250/ASSET/IMAGES/LARGE/ES1C01250_0005.JPEG

21. Pärnänen K, Karkman A, Tamminen M, Lyra C, Hultman J, Paulin L, et al. Evaluating the mobility potential of antibiotic resistance genes in environmental resistomes without metagenomics. Sci Reports 2016 61. 2016;6: 1–9. doi:10.1038/srep35790

22. Hindson BJ, Ness KD, Masquelier DA, Belgrader P, Heredia NJ, Makarewicz AJ, et al. High-throughput droplet digital PCR system for absolute quantitation of DNA copy number. Anal Chem. 2011;83: 8604–8610. doi:10.1021/ac202028g

23. Whale AS, Huggett JF, Tzonev S. Fundamentals of multiplexing with digital PCR. Biomol Detect Quantif. 2016;10: 15–23. doi:10.1016/J.BDQ.2016.05.002

24. Regan JF, Kamitaki N, Legler T, Cooper S, Klitgord N, Karlin-Neumann G, et al. A Rapid Molecular Approach for Chromosomal Phasing. PLoS One. 2015;10: e0118270. doi:10.1371/JOURNAL.PONE.0118270

25. Gillings MR, Gaze WH, Pruden A, Smalla K, Tiedje JM, Zhu YG. Using the class 1 integron-integrase gene as a proxy for anthropogenic pollution. ISME J. 2015;9: 1269–1279. doi:10.1038/ismej.2014.226

26. Partridge SR, Tsafnat G, Coiera E, Iredell JR. Gene cassettes and cassette arrays in mobile resistance integrons. FEMS Microbiol Rev. 2009;33: 757–784. doi:10.1111/J.1574-6976.2009.00175.X

27. Barraud O, Baclet MC, Denis F, Ploy MC. Quantitative multiplex real-time PCR for detecting class 1, 2 and 3 integrons. J Antimicrob Chemother. 2010;65: 1642–1645. doi:10.1093/JAC/DKQ167

28. Rocha J, Fernandes T, Riquelme M V., Zhu N, Pruden A, Manaia CM. Comparison of Culture- and Quantitative PCR-Based Indicators of Antibiotic Resistance in Wastewater, Recycled Water, and Tap Water. Int J Environ Res Public Heal 2019, Vol 16, Page 4217. 2019;16: 4217. doi:10.3390/IJERPH16214217

29. Land M, Hauser L, Jun SR, Nookaew I, Leuze MR, Ahn TH, et al. Insights from 20 years of bacterial genome sequencing. Funct Integr Genomics. 2015;15: 141. doi:10.1007/S10142-015-0433-4

30. Zhang AN, Li LG, Ma L, Gillings MR, Tiedje JM, Zhang T. Conserved phylogenetic distribution and limited antibiotic resistance of class 1 integrons revealed by assessing the bacterial genome and plasmid collection. Microbiome. 2018;6: 1–14. doi:10.1186/S40168-018-0516-2/FIGURES/5

31. Gweon HS, Shaw LP, Swann J, De Maio N, Abuoun M, Niehus R, et al. The impact of sequencing depth on the inferred taxonomic composition and AMR gene content of metagenomic samples. Environ Microbiomes. 2019;14: 7. doi:10.1186/s40793-019-0347-1

32. Link-Lenczowska D, Pallisgaard N, Cordua S, Zawada M, Czekalska S, Krochmalczyk D, et al. A comparison of qPCR and ddPCR used for quantification of the JAK2 V617F allele burden in Ph negative MPNs. Ann Hematol. 2018;97: 2299–2308. doi:10.1007/S00277-018-3451-1/TABLES/2

33. Alvarado A, Garcillán-Barcia MP, de la Cruz F. A degenerate primer MOB typing (DPMT) method to classify gamma-proteobacterial plasmids in clinical and environmental settings. PLoS One. 2012;7. doi:10.1371/journal.pone.0040438

34. Garcillán-Barcia Mp, Ruiz del Castillo B, Alvarado A, de la Cruz F, Martínez-Martínez L. Degenerate primer MOB typing of multiresistant clinical isolates of E. coli uncovers new plasmid backbones. Plasmid. 2015;77: 17–27. doi:10.1016/J.PLASMID.2014.11.003

35. Villa L, Carattoli A. Plasmid Typing and Classification. Methods Mol Biol. 2020;2075: 309– 321. doi:10.1007/978-1-4939-9877-7_22

36. Eastburn DJ, Huang Y, Pellegrino M, Sciambi A, Ptáček LJ, Abate AR. Microfluidic droplet enrichment for targeted sequencing. Nucleic Acids Res. 2015;43: e86–e86. doi:10.1093/NAR/GKV297

37. Dumke R, Barron M de la C, Oertel R, Helm B, Kallies R, Berendonk TU, et al. Evaluation of Two Methods to Concentrate SARS-CoV-2 from Untreated Wastewater. Pathog 2021, Vol 10, Page 195. 2021;10: 195. doi:10.3390/PATHOGENS10020195

38. Alygizakis N, Markou AN, Rousis NI, Galani A, Avgeris M, Adamopoulos PG, et al. Analytical methodologies for the detection of SARS-CoV-2 in wastewater: Protocols and future perspectives. TrAC Trends Anal Chem. 2021;134: 116125. doi:10.1016/J.TRAC.2020.116125

39. Pillay L, Amoah ID, Deepnarain N, Pillay K, Awolusi OO, Kumari S, et al. Monitoring changes in COVID-19 infection using wastewater-based epidemiology: A South African perspective. Sci Total Environ. 2021;786: 147273. doi:10.1016/J.SCITOTENV.2021.147273

